# Forest quality, forest area and the importance of beta-diversity for protecting Borneo’s beetle biodiversity

**DOI:** 10.1101/434134

**Authors:** Adam C. Sharp, Maxwell V. L. Barclay, Arthur Y. C. Chung, Guillaume de Rougemont, Edgar C. Turner, Robert M. Ewers

## Abstract

The lowland forest of Borneo is threatened by rapid logging for timber export and clearing for the expansion of timber and oil palm plantations. This combination of processes leaves behind landscapes dotted with small, often heavily-degraded forest fragments. The biodiversity value of such fragments, which are easily dismissed as worthless, is uncertain. We collected 187 taxa of rove beetles across a land-use gradient in Sabah, Malaysia, spanning pristine tropical lowland forest to heavily-degraded forest. Using these data, we identified shifts in alpha-, beta-, and gamma-diversity in response to forest quality and distance, then applied our findings from continuous expanses of forest to make predictions on hypothetical forest areas. We found that maintaining high forest quality is more important than forest area for conserving rare taxa (those important for conserving biodiversity per se), and that very small areas (10 ha) are likely to harbour the same richness of rove beetles as larger areas (100 ha) of equal forest quality. We estimate a decline in richness of 36% following heavy logging (removal of 95% of the vegetation biomass) from a forest area of 100 ha or less. Maintaining large forest area as well as high forest quality is important for conserving community composition, likely to be more important for conserving ecosystem functioning. We predict a decline of 35% in community diversity in conversion of a 100 ha area of unlogged forest to a 10 ha area of heavily-logged forest. Despite significant declines in alpha-diversity, beta-diversity within small rainforest areas may partially mitigate the loss of gamma-diversity, reinforcing the concept that beta-diversity is a dominant force determining the conservation of species in fragmented landscapes. In contrast to previous findings on larger animals, our results suggest that even small fragments of degraded forest might be important reservoirs of invertebrate biodiversity in tropical agriculture landscapes. These fragments, especially of lightly-logged forest, should be conserved where they occur and form an integral part of management for more sustainable agriculture in tropical landscapes.

## Introduction

Borneo’s forest is amongst the most biodiverse on the planet (Kier et al. 2005; de Bruyn et al. 2014), but that forest is threatened by rapid logging and clearing for the expansion of plantations (Fitzherbert et al. 2008; Lewis, Edwards & Galbraith 2015; Tsujino, Yumoto, Kitamura, Djamaluddin & Darnaedi 2016). Plantations support little biodiversity in comparison to adjacent natural habitats (Chung, Eggleton, Speight, Hammond & Chey 2000; Edwards et al. 2010; Fayle et al. 2010). While most studies agree that unlogged forest is irreplaceable for biodiversity (Gibson et al. 2011), the ongoing global expansion of logging and fragmentation (Taubert et al. 2018) requires us to also consider the value of degraded forest to conservation. This is especially important in the Malaysian Borneo state of Sabah, where large amounts of logged forest have been progressively added to the protected area network (Reynolds, Payne, Sinun, Mosigil & Walsh 2011).

Degraded forest can protect a large proportion of natural biodiversity (Edwards et al. 2011; Wearn et al. 2017). Modified forest is often highly variable in disturbance and structure (Burivalova, Sekercioglu & Koh 2014; Pfeifer et al. 2016), yet many studies on Borneo’s biodiversity have been unable to account for that variation and thus treat logged and unlogged as discrete forest types. Habitat quality needs to be considered on a continuous scale to understand the subtle responses of biodiversity to disturbance (Edwards et al. 2011; Burivalova, Sekercioglu & Koh 2014). The same is true for fragmentation - remnant forest fragments support a greater level of biodiversity than surrounding plantation (Edwards et al. 2010), and this biodiversity value increases with fragment size (Struebig et al. 2011). It is difficult, however, to directly quantify independent effects of forest quality and fragment size because the two are often interlinked (Tawatao et al. 2014; Taubert et al. 2018). Understanding how forest quality and area together impact biodiversity might highlight low-effort strategies for maintaining greater levels of natural diversity in agriculture-dominated landscapes.

To add further complexity to this knowledge-gap, diversity itself is comprised of three components that could all be impacted individually by forest degradation. Diversity at any one point is alpha-diversity and diversity between points is beta-diversity. Total diversity of an area is gamma-diversity (Whittaker 1972; Magurran 2004). Beta-diversity is often neglected from studies of logging and oil palm plantation impacts on biodiversity (but see Hamer & Hill 2000), yet is particularly relevant for a spatial problem like forest fragmentation (Tscharntke et al. 2012) and in highly diverse tropical forests where beta-diversity can be the largest contributor to gamma-diversity (Beck et al. 2012).

Previous work has examined the responses of beta-diversity to logging, and found that logged forest is highly beta-diverse (Hamer & Hill 2000; Pfeiffer & Mezger 2012; Kitching et al. 2013; Wearn et al. 2016). However, where beta-diversity has been quantified, it is often in metrics that are either directly dependent on alpha-diversity (Jost 2007) or dissimilar in scale (Benedick et al. 2006; Lucey et al. 2014). This not only prohibits the comparison of alpha-diversity with beta-diversity, but also invalidates calculation of beta over gradients in alpha (Jost 2007). We remain, therefore, largely uninformed regarding the independent response of beta-diversity to conversion of Borneo’s forest, despite the fact that beta-diversity is probably the largest contributor to forest-level gamma-diversity (Beck et al. 2012).

We aimed to quantify the independent impacts of forest quality and distance between sample sites on each component of biodiversity, paying close attention to how alpha- and beta-diversity shape gamma-diversity. To achieve this, we chose to study rove beetles (order: Coleoptera, family: Staphylinidae). Rove beetles were ideal for their high taxonomic and functional diversity (Barton, Gibb, Manning, Lindenmayer & Cunningham 2011), high abundance in tropical rainforests, and environmental sensitivity (Bohac 1999). While it is never possible to quantify biodiversity change in every taxon present, focussing an analysis on the rove beetles alone should provide conclusions of broad relevance to the wider ecosystem. We hypothesised that beta-diversity would make a larger contribution to the gamma-diversity of forest areas than alpha-diversity, and that beta-diversity would increase with distance and disturbance. We use our findings to predict diversity reductions that would result from different scenarios of forest degradation and conversion in Borneo. From these predictions, we generate rules of thumb that can be applied to inform management practice in preserving tropical biodiversity within predominantly agricultural landscapes.

## Methods

### STUDY AREA

This study was conducted at the Stability of Altered Forest Ecosystems (SAFE) Project (Ewers et al. 2011) in Sabah, Malaysian Borneo (Fig. 1). The project utilises a fractal sampling pattern that was designed specifically to study diversity at multiple spatial scales (Ewers et al. 2011; Marsh & Ewers 2013). The SAFE Project comprises 7,200 ha of forest that was unevenly logged once in the 1970’s (removing 113 m^3^ ha^−1^) and again between 2000 and 2008 (removing 66 m^3^ ha^−1^, Struebig et al. 2013), resulting in a logged forest area with widely varying forest quality (Pfeifer et al. 2016). Sample sites were spread across 10 discrete blocks (Ewers et al. 2011). One sampling block was in primary forest at Maliau Basin Conservation Area (MBCA), with two-thirds of sites in forest that has never been modified and one-third of sites in forest that has undergone light logging to obtain timber for construction of the adjacent field centre. A second sampling block was in a continuous expanse of logged forest. A third block was in a Virgin Jungle Reserve that had experience illegal logging around the periphery, and a further seven blocks were in or adjacent to the logged forest of the SAFE area (Fig. 1). Within each block, sample sites were clustered at three spatial scales. Three sites were separated at second-order scale by approximately 10^2.25^ m, with those clusters of sites separated at third-order scale by 10^2.75^ m and again at fourth-order scale by 10^3.25^ m. At the time of sampling, all locations were connected as part of a large, continuously forested area that extends across more than one million hectares (Reynolds, Payne, Sinun, Mosigil & Walsh 2011).

**Figure 1.**
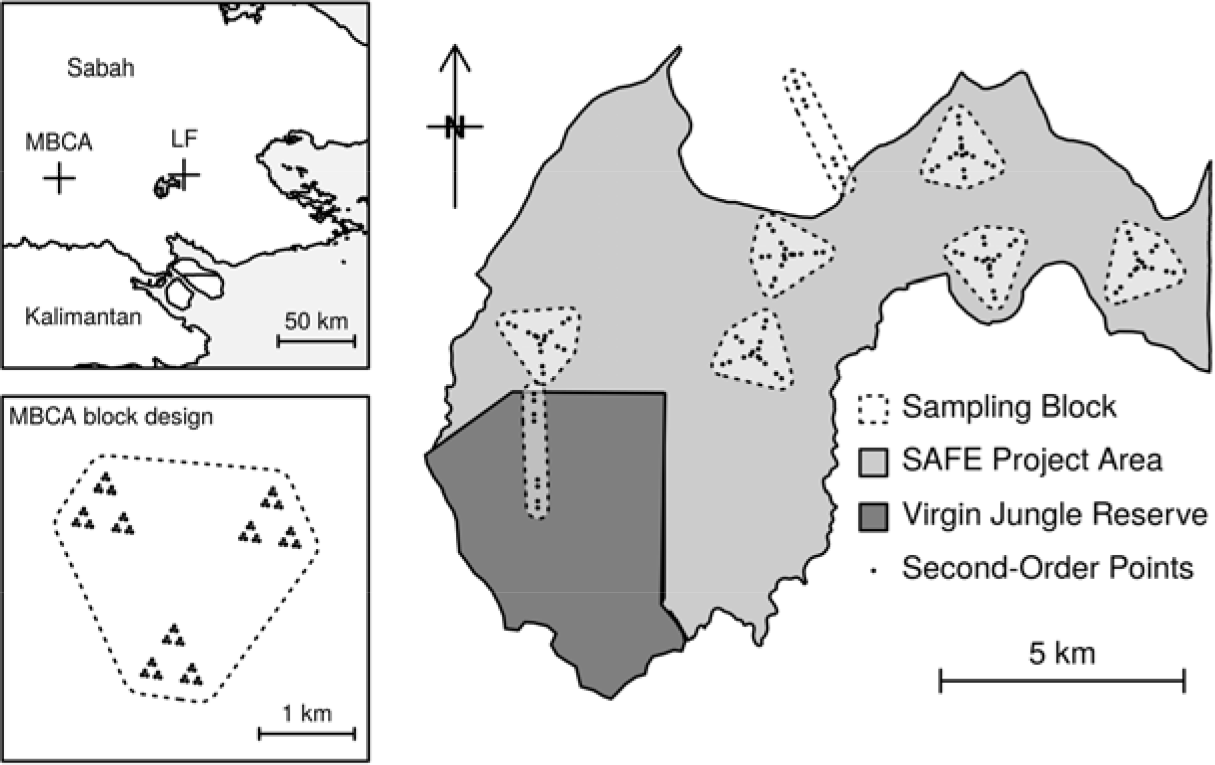
Arrangement of sampling blocks in Sabah, Malaysia. Sampling blocks outside of the SAFE Project area and VJR are indicated in the top-left panel by +. The sampling block design at MBCA is shown in the bottom-left panel and is of equivalent design to the logged forest control block labelled as LF (top-left).

### INSECT SAMPLING

We sampled insects at up to 166 second-order sites twice in 2011, once in February and again in November/December. Three insect traps were positioned around each site and were separated by 101.75 m (first-order scale). Traps were a design combining pitfall (diameter approximately 25 cm), flight-interception (surface area approximately 1 m^2^) and malaise traps and left active for three days. Insects were directed either upwards or downwards into collection bottles filled partially with 70% ethanol, and trap samples were combined at each site. Rove beetles were identified to the lowest taxonomic level possible, but the abundant subfamily Aleocharinae was removed from analysis as it proved impossible to identify them to any meaningful level.

### CALCULATING DIVERSITY METRICS

Most studies on forest conversion in Borneo employ species richness to examine diversity changes and are useful in assessing the vulnerability of threatened or charismatic taxa. However, taxa rarely coexist in equal numbers (Hubbell 2001) so relying solely on taxa richness to quantify diversity disregards community composition (measures of the identity and abundance of different taxa, Magurran 2004) which can change independently to species richness (Beck, Kitching & Linsenmair 2006; Banks-Leite, Ewers & Metzger 2012). It is important to consider both in biodiversity management to protect both rare or vulnerable taxa alongside the common taxa that likely contribute the majority of ecosystem function (Slade, Mann & Lewis 2011; Winfree, Fox, Williams, Reilly & Cariveau 2015). In recognition of this, we chose to calculate diversity using the equations derived by Jost (2007). These metrics generate independent measures of alpha- and beta-diversity, and allow the examination of trends in both taxa richness and community composition.

Each of alpha-, beta- and gamma-diversity were calculated in three ways: weighting in favour of rare taxa, weighting all taxa equally, and weighting in favour of common taxa, corresponding to q = 0, 1 and 2 respectively in Jost’s diversity equations. When q = 0, alpha-diversity is equivalent to mean taxa richness across a set of sample sites, and gamma-diversity is equivalent to total taxa richness for all sample sites combined. When q = 1, diversity measurements are equivalent to Shannon indices and, when q = 2, diversity measurements are equivalent to Simpson’s indices. These three weightings encompass a gradient of arguments for the conservation of biodiversity. A weighting of q = 0 treats all species equally, regardless of abundance, and is applicable to arguments based on conserving species richness and biodiversity *per se*. By contrast, q = 2 weights in favour of common species – those most likely to be dominating the ecology of the ecosystem (Slade, Mann & Lewis 2011; Winfree, Fox, Williams, Reilly & Cariveau 2015) – and is applicable to arguments about conserving ecosystem function. Only sites which caught at least one rove beetle were used, as some diversity metrics are undefined when gamma-diversity is 0 (Jost 2007).

### RELATING INSECT TRAP DATA TO FOREST DISTANCE

Examining the rove beetles caught at any one site would have limited our study to alpha-diversity alone. We chose to randomly group multiple sites together in order to develop estimates of beta-diversity and gamma-diversity. Distance between sites within a group was used as a proxy for forest area and related to each of beta- and gamma-diversity. Combinations of three sites were used as the simplest method of generating 2-dimensional shapes from second-order sites. All combinations were nested within blocks and sampling periods.

### DEFINING DISTANCE AND FOREST QUALITY VARIABLES

Each three-site combination was assigned continuous forest quality and between-site distance values. Quality was quantified by Pfeifer et al. (2016), who calculated estimates of above ground biomass (AGB) from 25 × 25 m (0.0625 ha) vegetation plots at each of the 166 second-order sites. High levels of disturbance were characterised by low AGB, while primary forest sites had the highest AGB. We estimated, from Pfeifer et al. (2016), that the mean unlogged above-ground biomass of vegetation was 524 ± 54 Mg/ha. We exploited the hierarchical nature of the fractal sampling design to assign a forest quality value to each three-site combination that was relevant to spatial scale. Combinations where the three second-order points were clustered around the same third-order point were given the mean AGB value of all second-order points clustered around that third-order point; the same approach was extended to the fourth-order. Three-site combinations over greater than fourth-order scale were given the mean AGB value of the entire sampling block. Distance between sites was calculated as the total centroid distance formed by the three chosen points (Fig. 2), and ranged from 400 m to 2,200 m.

**Figure 2.**
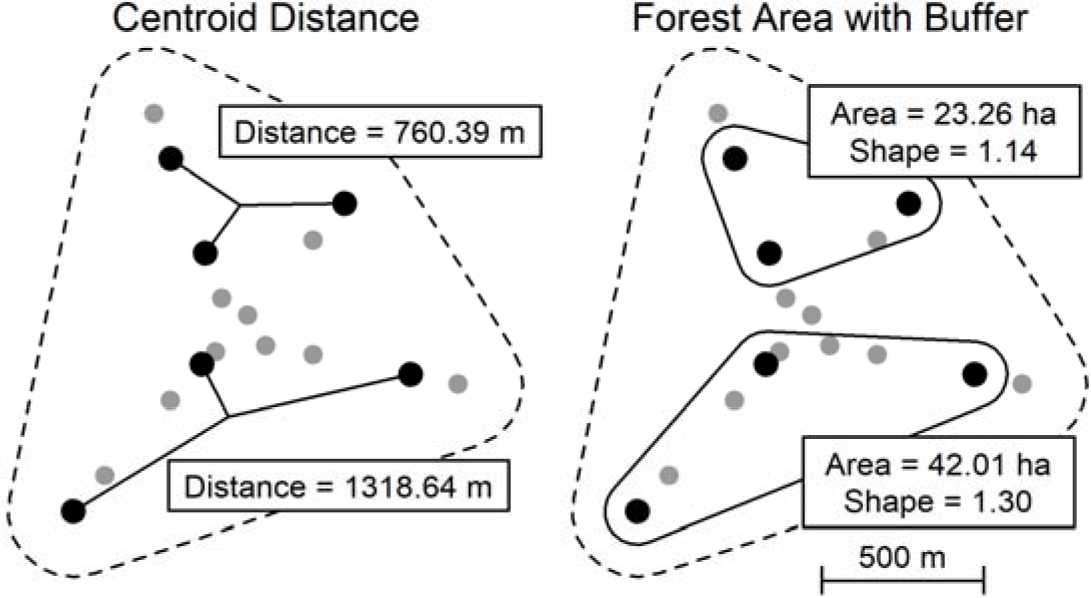
Random three-site combinations represented within a sampling block (dashed line). Sampling sites comprising one of the three sites within a single combination are black circles, while sampling sites which are not in a combination are grey circles.

### DEVELOPING BOOTSTRAP MODELS

To accurately model the response of rove beetle diversity to AGB and centroid distance, we had to group sites into combinations of three many times rather than just once. Within a classical linear model, this process would require resampling a single site to generate multiple data points, and therefore introduce a high level of pseudoreplication. We combatted this problem by randomly selecting 40 three-site groups in iteration, and fitting linear models predicting each of alpha-, beta- and gamma-diversity using q = 0, q = 1 and q = 2 within each iteration (nine models per iteration). All models except gamma-diversity where q = 0 were linear mixed models with log-transformed diversity as the response, AGB and distance and their interaction terms as fixed effects, and sampling period as a random intercept. The gamma-diversity/q = 0 models were generalized linear models with poisson error family, log-link function and an observation-level random intercept to account for overdispersion (Harrison 2014). All explanatory variables were log-transformed. Within any one iteration, no site was used more than once to avoid pseudoreplication, and we were able to sample from 6,684 possible three-site combinations. Overall model parameter estimates were taken as the mean of respective estimates from each iteration and were therefore effectively bootstrapped.

There was a weak positive correlation between sampling block area and AGB at the SAFE Project which manifested itself in false relationships between alpha-diversity and centroid distance. Alpha-diversity must be independent of distance or area (Whittaker 1972), and so we forced this independence on our bootstrapped dataset. This was achieved by randomly generating 100,000 iterations of 40 groups prior to analysis, fitting models of alpha-diversity within each of those iterations, and then selecting for analysis the 10,000 of those 100,000 iterations where the sum of squares of distance parameter estimates on alpha-diversity was minimised.

For each of the 10,000 iterations, model selection was achieved using backward selection of the terms AGB, AGB^2^, distance, and their interactions according to the rule of marginality. Applying Z-tests to determine whether mean parameter estimates differed significantly from 0 resulted in selection of all possible terms for beta- and gamma-diversity because of the large number of iterations we were able to perform (high number of iterations = low standard error and high Z-value). To increase the rigour of our model selection, we added the condition that effect sizes of each parameter estimate must be greater than 0.2 (a small effect according to Cohen, 1969, p.23).

### CREATING NOMINAL FOREST AREAS

To translate our results - generated from samples collected in continuous forest areas - to recommendations about the potential impact of forest fragmentation in agricultural landscapes, we extrapolated forest area from the three-site combinations (Fig. 2). We considered the area inside each three-site combination as being equivalent to the core of a forest fragment; i.e. the part of a forest fragment that is not influenced by the edge between forest and its surrounding matrix habitat. Work elsewhere has demonstrated that 100 m is the approximate extent to which forest edges influence diversity and abundance of tropical leaf-litter invertebrates (Laurance et al. 2002), and 80% of beetle species have edge effect extents of 100 m or less (Ewers & Didham 2008). We therefore added a 100 m buffer around the periphery of each three-site combination and refer to this total area as nominal forest area. Our three-site combinations ranged in nominal forest areas from approximately 10 ha to 100 ha. This represents an appropriate range for study because more than 90% of remnant fragment fragments following clearing of tropical forests are smaller than 100 ha (Ranta, Blom, Nimela, Joensuu & Siitonen 1998; Ribeiro, Metzger, Martensen, Ponzoni & Hirota 2009). The most recent trends in global deforestation show that forest fragments are ever-decreasing in area (Taubert et al. 2018).

We quantified the shape of each nominal forest area using the Shape Index (SI) derived by Patton (1975) and adapted for metric units (Didham & Ewers 2012). A value of SI = 1.0 represents a perfect circle, and greater values are progressively more elongate. Because the effects of forest fragment shape are likely dependent on the surrounding matrix, and because we were unable to simulate the complex shapes of fragments observed in real-world landscapes (Ewers & Didham 2008) using simple triangles, we chose to control for shape instead of including this variable in our models and so limited our selected forest areas to those with SI < 1.4.

Three further bootstrap models (one for each value of q) were fitted to predict gamma-diversity in the nominal forest areas from their size and forest quality. As before we generated 100,000 random iterations of 40 three-site combinations (from 4,593 possible combinations). We assigned each combination a value of AGB as before, but calculated the nominal forest area for each three-site combination instead of centroid distance. From these we selected the 10,000 iterations which minimised parameter estimates of nominal forest area on alpha-diversity, and fitted bootstrap models of gamma-diversity alone using the same methods to which we fitted our centroid distance models. We used these final gamma-diversity models to produce estimates of proportional change in total rove beetle diversity given proportional change in forest quality or nominal forest area.

## Results

A total of 11,352 rove beetles were caught over the two sampling periods, of which 2,905 belonged to subfamilies other than Aleocharinae. There were 1,138 beetles belonging to the Staphylininae, 1,083 to the Oxytelinae, 534 to the Paederinae, 87 to the Osoriinae, 37 to the Tachyporinae, 14 to the Euaesthetinae, six to the Omaliinae and six to the Steninae. From those eight subfamilies, there were 187 reproducible taxonomic units, of which 40 were named species accounting for 618 individuals. A further 143 unnamed taxa (n = 1,674) were known to be separate species of non-aleocharine rove beetle. The remaining 613 individuals were identifiable to one out of the genera *Anotylus* (n = 315), *Mitomorphus* (n = 82) or *Thinocharis* (n = 71) or the tribe Xantholini (n = 145). A total of 90 sites caught non-aleocharine rove beetles in the first sampling period and 144 in the second. The number of non-aleocharine rove beetles caught per site increased with AGB (Fig. 3).

**Figure 3.**
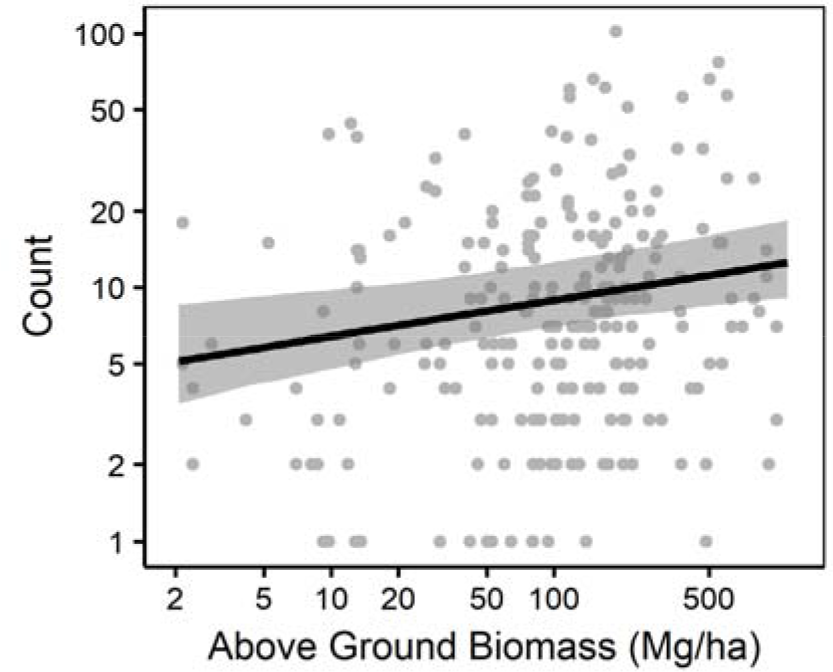
Abundance of non-aleocharine rove beetles caught at individual sites across the land-use gradient. Model fit is from a generalized mixed model with poisson error, log link function, random intercept for sampling period and second observation-level random intercept for modelling overdispersion (Harrison 2014). Grey area represents 95% confidence intervals, which were estimated via bootstrapping.

Where q = 0 (taxa richness, Table 1), alpha-diversity at the lowest measured AGB was approximately 30% of that at primary forest AGB (524 Mg/ha). There was a peak in alpha-diversity at around 200 Mg/ha; equivalent to the highest values of AGB measured in logged forest (Fig. 4a). The bootstrap beta-diversity model for q = 0 (pseudo-R^2^ = 0.44) explained more variation than alpha-diversity (pseudo-R^2^ = 0.09). Beta-diversity was highest at low AGB and was greater than alpha-diversity in heavily-logged forest (AGB < 57 Mg/ha). There was a small interaction effect between AGB and centroid distance whereby beta-diversity increased with distance at highest forest quality (Fig. 4d). Th effect of distance was small enough that it was not selected in our models of gamma-diversity where q = 0, and only AGB influenced gamma-diversity (Fig. 4g). As with alpha-diversity, gamma-diversity was greatest at around 200 Mg/ha.

**Table 1.** Parameter estimates for bootstrap models predicting each of alpha-, beta- and gamma- diversity from AGB and centroid distance. Only final models after model selection are shown. Pseudo-R^2^ values are the mean of McFadden’s (1974) pseudo-R^2^ when the selected model was applied to each of 10,000 iterations of data combinations. Mean, Standard Deviation (SD) and Standard Error (SE) are those of the respective parameter from 10,000 fitted mixed models. The effect size, d, is the absolute value of Mean/S.D. (Cohen 1969).

| Weighting | Component | Parameter | Mean | SD | d | SE | Z | P |
| --- | --- | --- | --- | --- | --- | --- | --- | --- |
| q = 0 | Alpha | Intercept | -9.95 | 3.45 | 2.88 | 0.03 | -288.14 | < 0.001 |
| | Pseudo-$R^2$ = 0.09 | AGB | 9.94 | 3.09 | 3.21 | 0.03 | 321.20 | < 0.001 |
|  |  | AGB<sup>2</sup> | -2.13 | 0.68 | 3.14 | 0.01 | -314.26 | < 0.001 |
|  | Beta | Intercept | -2.65 | 42.23 | 0.06 | 0.42 | -6.28 | < 0.001 |
| | Pseudo-$R^2$ = 0.44 | AGB | 5.55 | 37.70 | 0.15 | 0.38 | 14.73 | < 0.001 |
|  |  | AGB<sup>2</sup> | -1.51 | 8.27 | 0.18 | 0.08 | -18.27 | < 0.001 |
|  |  | Distance | 2.59 | 13.99 | 0.19 | 0.14 | 18.51 | < 0.001 |
|  |  | AGB x Distance | -2.50 | 12.48 | 0.20 | 0.12 | -20.03 | < 0.001 |
|  |  | AGB<sup>2</sup> x Distance | 0.60 | 2.74 | 0.22 | 0.03 | 22.07 | < 0.001 |
|  | Gamma | Intercept | -7.06 | 3.07 | 2.30 | 0.03 | -230.22 | < 0.001 |
| | Pseudo-$R^2$ = 0.05 | AGB | 8.44 | 2.76 | 3.06 | 0.03 | 306.15 | < 0.001 |
|  |  | AGB<sup>2</sup> | -1.87 | 0.61 | 3.08 | 0.01 | -307.67 | < 0.001 |
| q = 1 | Alpha | Intercept | -7.22 | 2.48 | 2.91 | 0.02 | -290.52 | < 0.001 |
| | Pseudo-$R^2$ = 0.11 | AGB | 7.35 | 2.25 | 3.27 | 0.02 | 327.05 | < 0.001 |
|  |  | AGB<sup>2</sup> | -1.59 | 0.50 | 3.20 | 0.00 | -320.27 | < 0.001 |
|  | Beta | Intercept | 4.54 | 3.16 | 1.43 | 0.03 | 143.48 | < 0.001 |

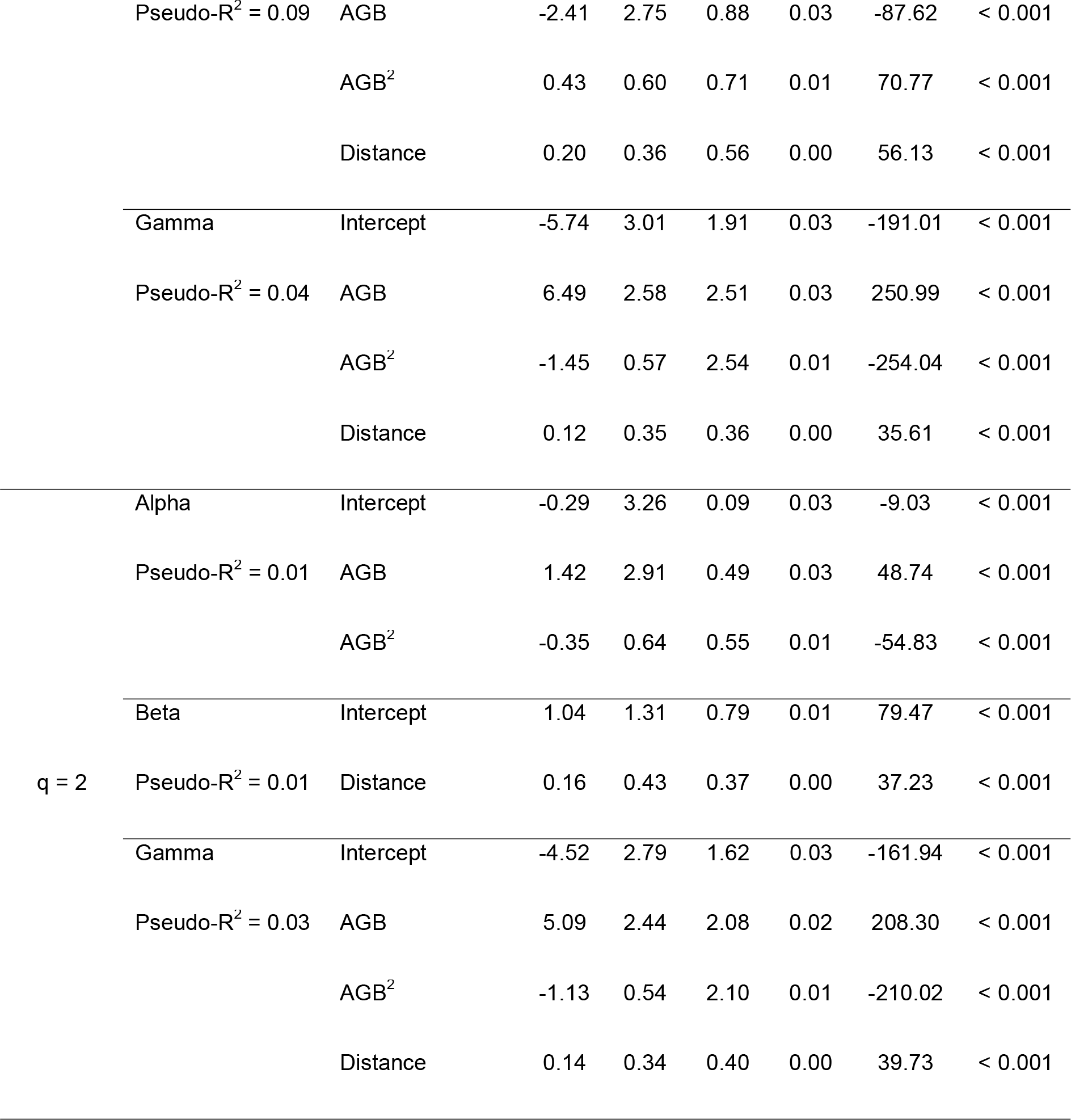

**Figure 4.**
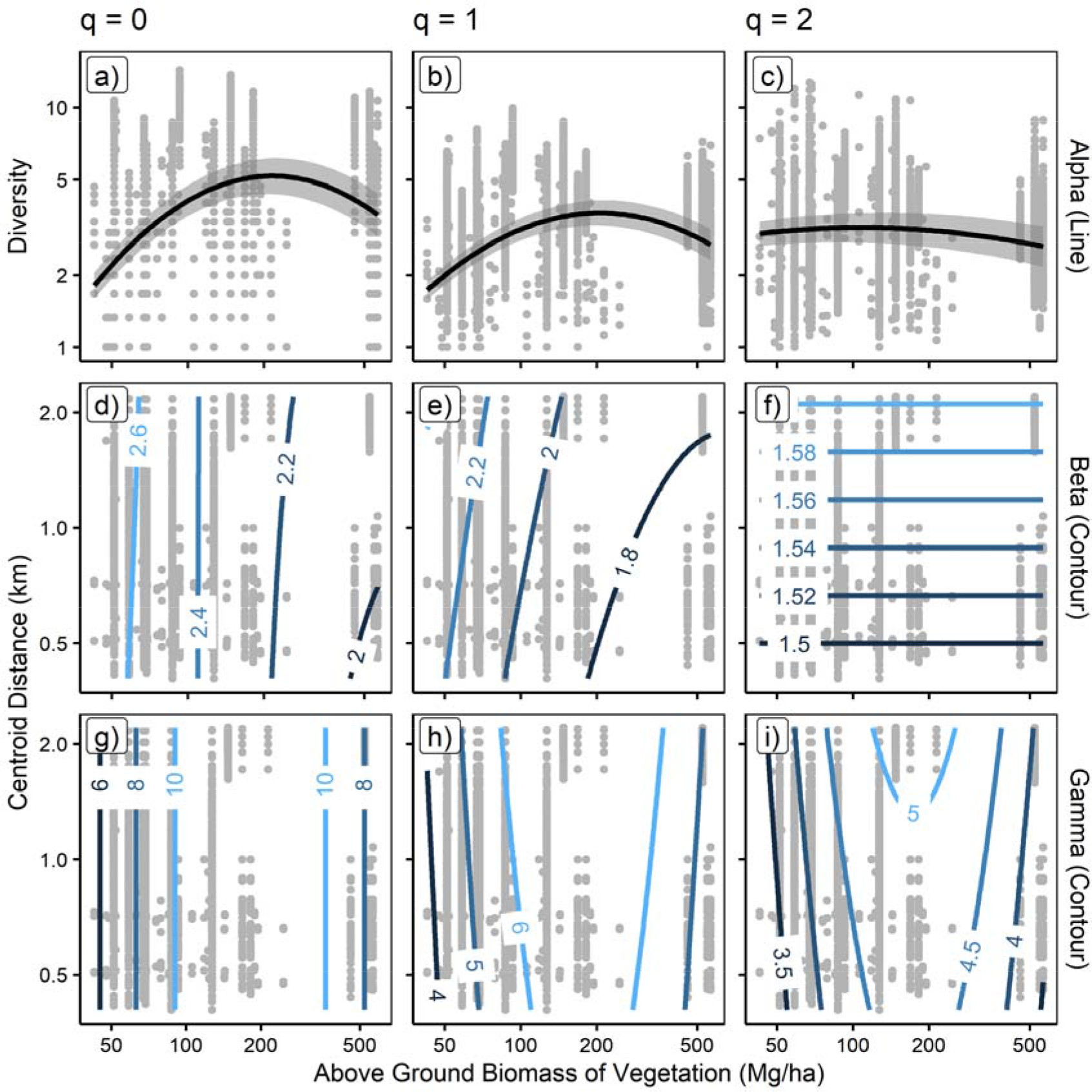
Responses of each of alpha-, beta- and gamma-diversity to AGB and centroid distance (beta- and gamma-diversity only). Alpha-diversity is represented as a scatter plot with no relation to centroid distance. Model estimate for alpha-diversity is represented by a black line, with 95% confidence intervals represented in grey. Beta- and gamma-diversity are both represented as contour plots as they can respond to both AGB and centroid distance. All axes are log-transformed.

AGB had a similar effect size on alpha-diversity where q = 1 (individual-weighted diversity, Table 1) compared to q = 0, and peaked around 200 Mg/ha (Fig. 4b). Alpha-diversity for q = 1 at lowest AGB was around 41% of that of primary forest AGB. Our beta-diversity model for q =1 explained approximately the same amount of variation as the alpha-diversity model (pseudo-R^2^ = 0.09 and 0.11 respectively). Beta-diversity increased with centroid distance across all values of AGB (albeit a small effect) and was highest at low AGB and high distance (Fig. 4e). Gamma-diversity where q = 1 increased with both AGB and centroid distance, with AGB having a far stronger effect, and was highest again at intermediate forest quality (Fig. 4h).

The models of diversity where q = 2 explained very little variation in the data (Table 1). We detected very small shifts in alpha-diversity, which was highest in logged forest (Fig. 4c), and beta-diversity, which increased slowly with centroid distance (Fig. 4f). As a result, gamma-diversity for q = 2 peaked in logged forest and increased with centroid distance (Fig. 4i), but the effect of AGB was stronger than that of distance.

We estimated proportional loss of gamma-diversity for each of q = 0, q = 1 and q = 2 in response to proportional losses in vegetation biomass and nominal forest area (Table 2). Where q = 0 (taxa richness), gamma-diversity after heavy logging (95% AGB removed) was reduced by 36% compared to gamma-diversity in an unlogged 100 ha forest area (Fig. 5a). Gamma-diversity for taxa richness also peaked in lightly-logged forest (66% AGB removed) at 164%. Nominal forest area had no effect on total taxa richness. Forest area did affect gamma-diversity where abundance was considered (q = 1 and 2), but this effect was smaller than that of AGB (Fig. 5b-c). Reducing a 100 ha nominal forest area to a 10 ha area while maintaining forest quality would result in a loss of 10% of the q = 1 gamma-diversity and 9% of the q = 2 gamma-diversity. Combining this reduction in forest area with heavy logging (removal of 95% of AGB/ha) would result in a 35% loss of gamma-diversity where q = 1 and 31% loss of gamma-diversity where q = 2.

**Table 2.** Parameter estimates for bootstrap models predicting gamma-diversity from AGB and nominal forest area. Only final models after model selection are shown. Values are as in Table 1.

| Weighting | Parameter | Mean | SD | d | SE | Z | P |
| --- | --- | --- | --- | --- | --- | --- | --- |
| q = 0 | Intercept | -8.12 | 3.19 | 2.54 | 0.03 | -254.45 | < 0.001 |
| Pseudo-R<sup>2</sup> = 0.06 | AGB | 9.42 | 2.89 | 3.27 | 0.03 | 326.50 | < 0.001 |
|  | AGB<sup>2</sup> | -2.09 | 0.64 | 3.27 | 0.01 | -327.30 | < 0.001 |
| q = 1 | Intercept | -6.66 | 3.01 | 2.21 | 0.03 | -221.04 | < 0.001 |
| Pseudo-R<sup>2</sup> = 0.06 | AGB | 7.57 | 2.72 | 2.78 | 0.03 | 277.98 | < 0.001 |
|  | AGB<sup>2</sup> | -1.69 | 0.60 | 2.81 | 0.01 | -281.09 | < 0.001 |
|  | Area | 0.08 | 0.28 | 0.28 | 0.00 | 27.99 | < 0.001 |
| q = 2 | Intercept | -5.23 | 2.81 | 1.86 | 0.03 | -185.92 | < 0.001 |
| Pseudo-R<sup>2</sup> = 0.04 | AGB | 6.03 | 2.57 | 2.35 | 0.03 | 235.02 | < 0.001 |
|  | AGB<sup>2</sup> | -1.34 | 0.57 | 2.37 | 0.01 | -236.96 | < 0.001 |
|  | Area | 0.07 | 0.28 | 0.24 | 0.00 | 23.93 | < 0.001 |

**Figure 5.**
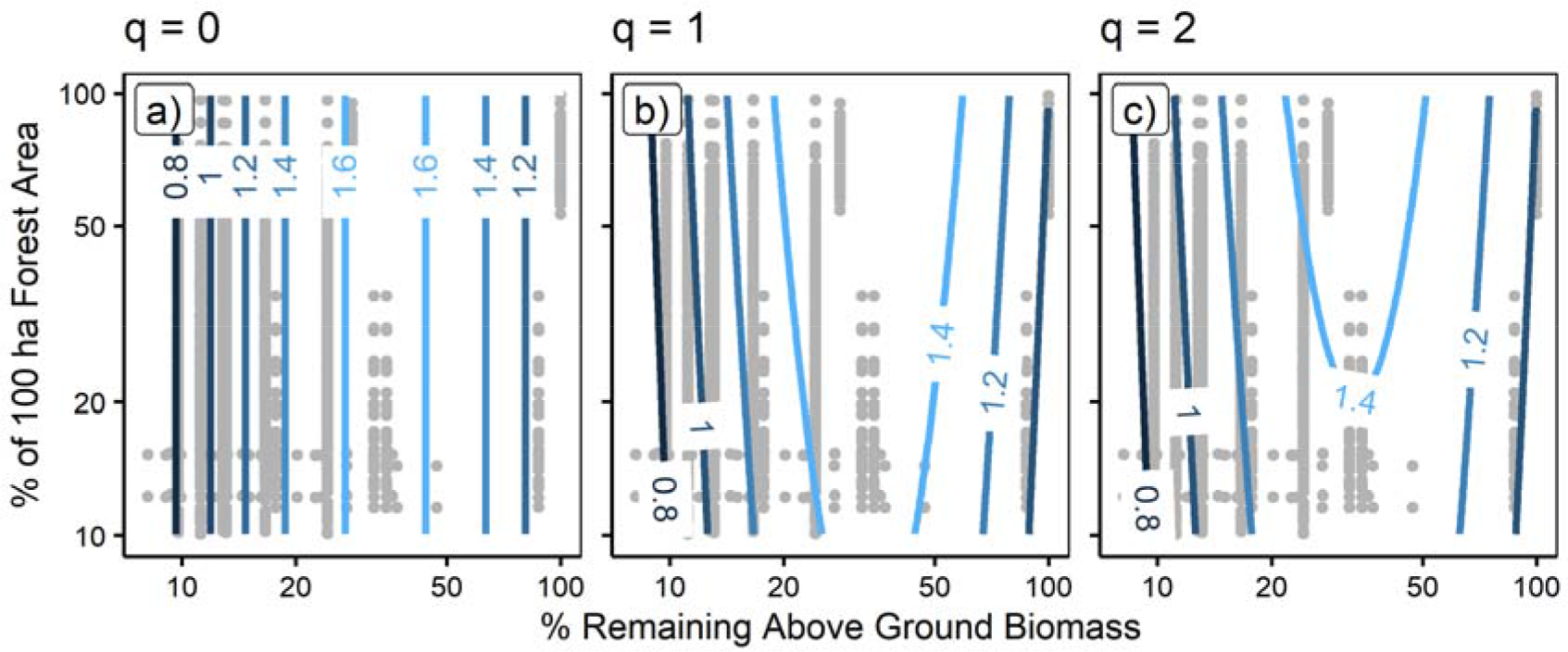
Proportional responses of gamma-diversity to percentage changes in forest quality and forest area (starting from 100 ha). Model estimates are represented as contours.

## Discussion

Preserving both high forest quality and protecting as much forest area as possible are both important to the conservation of biodiversity in Borneo. Our data for rove beetles – an abundant and ecologically important group of invertebrates (Bohac 1999) – demonstrate strongly contrasting responses to forest quality and forest area at the spatial scales relevant to heavily fragmented landscapes, with clear implications for forest management in the region. Our data indicate that forest quality is more important than area in propagating biodiversity at these small scales, but even the smallest, most-degraded forest areas may maintain a relatively large amount of gamma-diversity through shifts in beta-diversity.

In unlogged forest, high taxa richness (q = 0) is a result of there being many taxa at any given point (high alpha-diversity). In a heterogeneous, unlogged landscape, increasing sample area is likely to encompass a greater array of habitat types and their associated specialist taxa (Báldi 2008), explaining the small increase in richness beta-diversity with distance in high-quality forest. This did not, however, translate into an increase in gamma-diversity with nominal forest area. Other studies on forest fragments in SE Asia have found that larger fragments harboured greater invertebrate richness (Benedick et al. 2006; Lucey et al. 2014), but these analyses all contained data extending across a much greater range of fragment areas (over 123,000 ha and 500 ha respectively) than our largest nominal forest area (100 ha). It is possible that the spatial scale of forest areas we examined was not large enough to detect area-mediated shifts in gamma-diversity where q = 0. Reconciling previous results with ours would suggest that fragments greater in size than our maximum nominal forest area (100 ha) have large increases in within-fragment beta-diversity that go beyond the area-related increases we have observed in relatively small areas. However, our results suggest that in forest areas of < 100 ha, a size range that encompasses more than 90% of tropical forest fragments (Ranta, Blom, Nimela, Joensuu & Siitonen 1998; Ribeiro, Metzger, Martensen, Ponzoni & Hirota 2009), habitat quality is a far stronger predictor of taxa richness than area.

Our results indicate that in highly-disturbed forest, beta-diversity in taxa richness (Fig. 4d) mitigates substantial losses in alpha-diversity (Fig.4a) to maintain moderately high total richness (Fig. 4g). Beta-diversity is high for even small, heavily-logged areas, as we hypothesised. There are greater point-to-point differences in the taxa comprising rove beetle communities in such forest, not because the community is more spatially heterogeneous, but because naturally-common taxa have become less abundant (Fig. 3) and are therefore observed less frequently. The importance of beta-diversity to maintenance of high species richness in disturbed landscapes has been demonstrated previously in Borneo. For example, Wearn et al. (2016) showed that high beta-diversity contributes to the high richness of small and large mammals in Bornean logged forest, and Wang & Foster (2015) confirmed that high spatial beta-diversity can serve to maintain gamma-diversity of ants in plantation compared to natural forest. Our results therefore support past findings that beta-diversity plays a central role in mitigating biodiversity losses in human-modified landscapes (Tscharntke et al. 2012).

When all individuals of all species were equally weighted (q = 1), we found the highest alpha-diversity in unlogged and lightly-logged forest (Fig. 4b), indicating that all taxa in the community have relatively equal abundance at any given point. By contrast, lower alpha-diversity in highly-disturbed forest indicates that the community is dominated by a smaller number of taxa that make up a greater proportion of individuals at any one point. The high beta-diversity accompanying low alpha-diversity (Fig. 4e) suggests that the dominant taxa varies among sample points. The increase in beta-diversity with distance indicates that for any level of forest quality, large areas should be preserved to protect a community in which no single taxon dominates the entire area. This effect of this relationship on gamma-diversity was small at the spatial scales we examined, but may become more important at larger spatial scales (Benedick et al. 2006; Lucey et al. 2014).

Our models explained very little of the variation in the data where we gave higher weighting to the most common taxa (q = 2). As a result, we conclude that in comparison to rare taxa, there is little change in the diversity of common rove beetle taxa in response to forest quality and forest area. The apparent resilience of these common taxa to forest modification suggests that they, even in unlogged forest, are relatively generalist and are remarkably tolerant of any changes that might have accompanied the forest disturbance. It is these taxa which are most likely responsible for a large proportion of the ecosystem functioning in the landscape (Slade, Mann & Lewis 2011; Winfree, Fox, Williams, Reilly & Cariveau 2015), and our results therefore support previous reports that ecological processes have strong resilience to logging (Gray, Slade, Mann & Lewis 2014; Ewers et al. 2015).

All our models of gamma-diversity predicted peaks in overall diversity in lightly-logged forest (Fig. 5) due to high alpha-diversity (Fig. 4a-c). While unlogged forest is largely accepted to be more diverse than logged forest at greater scales (Gibson et al. 2011), our results suggest that this may not be true at these very small spatial scales. From an ecological point of view, these findings warn against studying “logged” forest as a discrete habitat category instead of a highly-variable array of mix of forest types (Burivalova, Sekercioglu & Koh 2014; Pfeifer et al. 2016). From an applied perspective, these findings suggest that preserving small areas of lightly logged forest may be a relatively low-effort, highly-effective method of boosting biodiversity in agriculture-dominated landscapes.

Our approach of simulating nominal forest areas has several advantages over using data from existing forest fragments, but is not perfect. Small fragments tend to be more heavily disturbed (Tawatao et al. 2014), which would have prevented us from separating the independent and interactive effects of fragment area and forest quality in real-world fragments. The real-world size and shape of forest fragments also confounds the explicit examination of beta-diversity. Moreover, using sub-samples from a large dataset collected in continuous habitats allowed us to make predictions for far more fragments than could possibly be sampled in fragmented landscapes. However, because our data did not come from existing fragments, our predictions about the exact effect of forest area on diversity retain a degree of uncertainty. We were not able to examine habitat shape because our sampling design was limited to very simple triangles, and so are unable to make generalisations to the convoluted shapes of real-world forest fragments (Ranta, Blom, Nimela, Joensuu & Siitonen 1998; Ewers & Didham 2008). Our design was also not able to identify changes in the invertebrate community which are linked to habitat edges (Ewers & Didham 2008; Terraube et al. 2016), a key contributor to shape effects.

Despite these limitations, our predicted losses in taxa richness were roughly equivalent to those observed in studies of real-world fragmentation scenarios in the same geographic region. Tawatao et al. (2014) found that small, heavily-degraded forest fragments had a 50% reduction in leaf-litter ant richness (fragment gamma-diversity) compared to larger, high-quality forest fragments. This percentage is close to our estimated 36% decrease in rove beetle taxa richness under the same conditions. Similarly, Didham, Hammond, Lawton, Eggleton & Stork (1998) observed a loss of 19% of common beetle species comparing Central Amazonian forest fragments across the same range of areas we were able to examine. This finding appears to concur with our conclusion that the common species are, although still sensitive to some extent to forest quality and area, largely resilient to forest disturbance. We argue that despite the short-falls in our method of modelling nominal forest areas, our approach represents a pragmatic solution to informing managers in how best to mitigate biodiversity losses in heavily-fragmented landscapes.

We found that conserving rare taxa of non-aleocharine rove beetles will require high quality forest to be maintained (Fig. 5a). Simultaneously, conserving common taxa, and by extension safeguarding the functioning of degraded forest ecosystems, will require retention of large forest fragments (Fig. 5b-c). We detected relatively minor declines in diversity in even the most degraded scenarios (Fig. 5), with evidence that increasing beta-diversity plays an important role in offsetting declines in alpha-diversity (Fig. 4d-e). Overall, our results indicate the balance between changes to alpha- and beta-diversity results in a remarkable biodiversity value for what, at first glance, might be considered near-worthless habitat: heavily-degraded forest.

## Acknowledgements

We thank the Royal Society’s South East Asia Rainforest Research Partnership for logistical support, and the Sabah Foundation, Maliau Basin Management Committee, the State Secretary, Sabah Chief Minister’s Departments, the Malaysian Economic Planning Unit and the Sabah Biodiversity Council for permission to conduct research. Data in the field were largely collected with the help and support of SAFE Project’s team of Research Assistants. This work was funded by the Sime Darby Foundation. This paper is a contribution to Imperial College's Grand Challenges in Ecosystems and the Environment initiative.

